# Connectome-scale self-supervised representation learning reveals neuronal organization beyond canonical labels

**DOI:** 10.64898/2026.06.30.735468

**Authors:** Te Shi, Yinda Chen, Chunxiuzi Liu, Ruobing Zhang

## Abstract

Dense electron-microscopy connectomes provide synaptic-resolution maps of neuronal structure and wiring, but learning scalable representations that integrate structure and connectivity for connectome discovery with minimal human intervention remains difficult. Here we present a self-supervised framework for structure–connectivity representation learning in dense connectomes. A hierarchical graph neural network with skeleton decomposition enables contrastive learning from finely sampled FlyWire neuronal skeletons, showing that fine skeletons preserve substantially richer identity information than coarse representations. Coordinate-free topology reduces developmental and geometric confounds, improving clustering and label-efficient inference. We then use learned structural embeddings as continuous descriptors of synaptic partners to construct structure-driven connectivity representations, improving subtype discrimination without predefined partner-type labels. Iterative multi-hop learning further reveals higher-order organization, including hemispheric connectivity lateralization and connectivity-defined subgroups. Attention analysis links these differences to specific synaptic partners. Together, these results establish a self-supervised and scalable framework for discovering neuronal identity and connectome organization in a large-scale dense connectome.

## Main

Understanding neural computation requires compact, informative representations of neuronal structure and wiring. Recent advances in serial-section electron microscopy have produced dense connectomes with synaptic-resolution reconstructions for up to hundreds of thousands of neurons, including the Drosophila hemibrain, FlyWire, and MICrONS datasets [1–3]. These resources provide an unprecedented opportunity to study neuronal identity, circuit organization and structure–function relationships at scale, but they also create a fundamental computational challenge: how to convert richly detailed neuronal reconstructions into compact, informative and transferable representations that support both cell-level and circuit-level analysis with minimal manual effort.

Existing approaches only partially address this challenge. After neuronal morphologies have been reconstructed, downstream connectomic analyses still often depend on predefined neuron types, curated partner labels, expert-guided clustering or iterative refinement [4–9]. Such workflows can be powerful in well-studied systems, but they are difficult to scale to newly reconstructed or less extensively annotated connectomes, where the informative features are not known in advance and manual analysis remains expensive. This creates a need for representation-learning methods that can extract neuronal organization directly from reconstructed skeletons and synapses, while reducing reliance on curated labels and expert-defined priors.

Recent work has begun to apply representation learning and graph neural networks (GNNs) to connectomic data [10, 11]. However, there are still several key obstacles. One is that biologically informative distinctions often arise at fine structural scales, where synapse placement, local branching patterns, and the spatial distribution of inputs and outputs along neurites can distinguish neuronal types that appear similar under coarse skeletonization [3, 5, 12]. Ideally, representation learning would operate directly on finely sampled neuronal trees with mapped synaptic sites. In practice, complete neuronal skeletons can contain thousands of nodes, making direct graph learning memory-intensive and difficult to scale for contrastive training [11, 13– 16]. To overcome this bottleneck, we introduce a hierarchical skeleton decomposition strategy together with a hierarchical graph neural network architecture. The framework decomposes each neuronal tree into smaller subgraphs, encodes local structure, and recursively integrates these embeddings into whole-neuron representations, enabling self-supervised contrastive learning on high-resolution skeletons under practical memory constraints.

Beyond scalability, dense connectomes raise a second representational challenge: absolute anatomical coordinates can both enrich and confound neuronal representations. Coordinate-aware morphology captures spatial embedding and fine geometric detail, but neurons of the same type may occupy mirrored hemispheres, translated positions or curved anatomical surfaces. In such cases, embeddings may reflect developmental variation, hemispheric placement or positional symmetry rather than intrinsic neuronal identity. We therefore distinguish coordinate-aware morphology from coordinate-free topology. In the topology regime, explicit spatial coordinates are removed while skeletal structure, branching organization and node-level synaptic annotations are preserved, allowing us to test whether topology provides a more robust representation layer for clustering and label-efficient neuron typing.

Finally, a third limitation is that connectomic identity is defined not only by a neuron’s intrinsic structure, but also by the partner neurons with which it forms synapses. However, many existing approaches treat morphology and connectivity as separate sources of information [17–19], limiting their ability to capture a holistic representation of connectomic organization. To address this bottleneck, we introduce a structure-driven connectivity representation that derives connectivity features from learned neuronal structure embeddings rather than from manually defined partner labels. Each neuron is first encoded by its structural representation learned from the skeleton. For a target neuron, each synaptic site is then assigned the learned embedding of its synaptic partner, so that partner-neuron identity is represented as a continuous vector at the level of individual synapses. A graph neural network with attention block then aggregates these partner embeddings across the target skeleton to produce a connectivity-aware neuronal representation. In this way, structural and connectivity information are integrated within a single learning framework, enabling connectivity-aware analysis without requiring predefined partner-type labels.

In this work, we present an annotation-light framework for learning integrated structure–connectivity representations in dense connectomes. Applied to FlyWire (“FlyWire Brain Dataset”), the framework enables self-supervised contrastive learning on finely sampled skeletons with 3,500 nodes, improves neuronal classification over coarse structural representations, and separates coordinate-aware morphology from coordinate-free topology. We show that topology provides a more robust representation layer for clustering and label-efficient neuron typing by reducing sensitivity to developmental variation, spatial position and anatomical symmetry. We further show that structure-driven connectivity enhances neuronal identity representations and reveals identity-relevant connections without requiring predefined partner-type labels. Together, these results establish a scalable strategy for downstream connectome analysis with reduced dependence on curated labels and expert-defined priors.

### Hierarchical graph learning enables representation learning from finely sampled skeletons

As a starting point, we aimed for structural precision sufficient to fully leverage the fine-scale neuronal morphology revealed by electron microscopy. To achieve this, we analyzed the meaningful nodes in FlyWire skeletons—nodes that are closely associated with synaptic connectivity or that play a critical role in the neuron’s topological structure, such as branch or terminal nodes (Fig. 1a–c). Based on the cumulative distribution of meaningful node counts, we selected 3,500 nodes as a practical upper bound, capturing 96.37% of neurons in the dataset.

**Figure 1.**
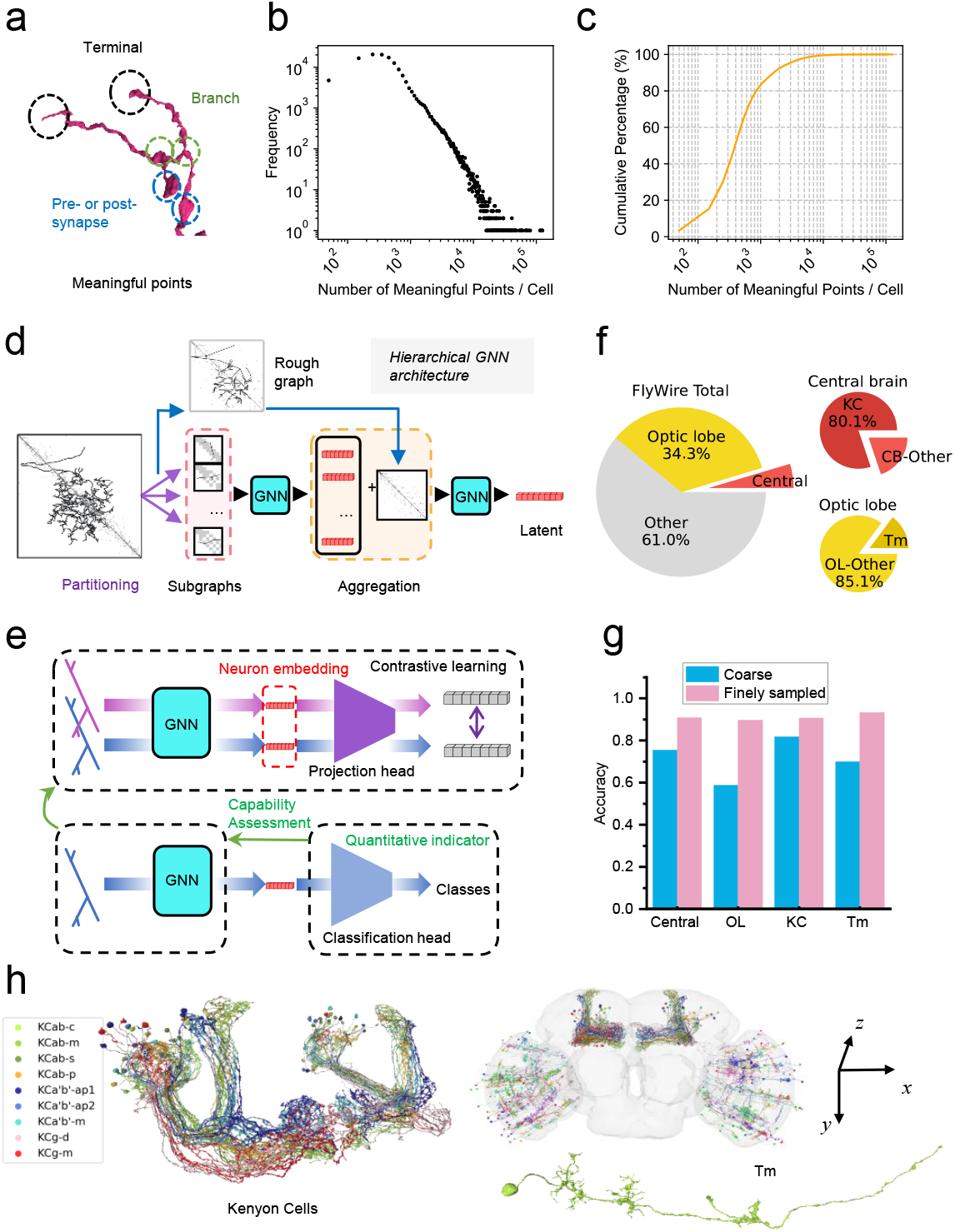
(a) Example of a neuron with meaningful points (points with terminal, branch, pre- or post-synapse labels). (b) Distribution of meaningful node counts across all neuron skeletons in FlyWire. (c) Cumulative percentage of neuron skeletons as a function of meaningful node counts. (d) Schematic of the hierarchical GNN architecture. The input graph is partitioned into subgraphs, locally encoded by a GNN, and subsequently aggregated by another GNN based on inter-subgraph adjacency. (e) Schematic of the contrastive learning and classification tasks. Classification performance is used as a quantitative indicator to evaluate the quality of learned representations under different input modalities. (f) Pie graph of the target neurons groups of four tasks. (g) Classification accuracy improves from coarse to finely sampled skeletons across four tasks: Central Brain (CB), Optic Lobe (OL), Kenyon Cells (KC), and Transmedullary neurons (Tm). (h) Example morphologies of KC and Tm neurons.

To handle skeletons at this scale, we developed a skeleton decomposition strategy together with a hierarchical graph neural network architecture (HGNNA, Fig. 1d). Rather than applying a single GNN to a graph containing thousands of nodes, the framework partitions each neuronal tree into smaller subgraphs and processes them hierarchically. Local structure is encoded first, and these local embeddings are then recursively integrated into higher-level representations. This makes graph contrastive learning on fine, synapse-aware skeletons computationally feasible.

We next asked whether the additional information preserved in fine skeletons improves downstream neuronal classification. After contrastive pretraining, the projection head was replaced with a classification head (Fig. 1e) and evaluated on four FlyWire tasks: central brain (CB), optic lobe (OL), Kenyon cells (KC), and transmedullary neurons (Tm). These tasks span both broad neuronal populations and more specific subtype distinctions. CB and OL together constitute 38.9% of all neurons in FlyWire and represent broad central and peripheral neuronal populations, respectively (Fig. 1f). KC and Tm are more specific neuronal classes within CB and OL (Fig. 1f), and can each be subdivided into finer types, with representative morphologies and spatial distributions shown in Fig. 1h.

Finely sampled skeletons consistently outperformed coarse skeletons across morphology-based classification on four neuronal lineages. Accuracy rose by +15.4 percentage points (75.4%→90.8%) in CB, +31.0 pp (58.7%→89.7%) in OL, +9.0 pp (81.7%→90.8%) in KC, and +23.2 pp (70.0%→93.2%) in Tm. These gains indicate that fine structural sampling is not a cosmetic refinement: it supplies substantially richer morphological information for deep-learning model across major compartments of the FlyWire connectome.

### Topological representations of neurons are robust to geometric and developmental confounds that bias coordinate-aware morphology

While fine-grained morphological representations demonstrate strong expressive power for neuronal classification, they can be confounded by functionally irrelevant developmental structure and spatial symmetry across neuron classes [20]. As illustrated in Fig. 2a, coordinate-aware morphology encodes explicit geometric information that may introduce spurious variation, thereby limiting its scalability.

**Figure 2.**
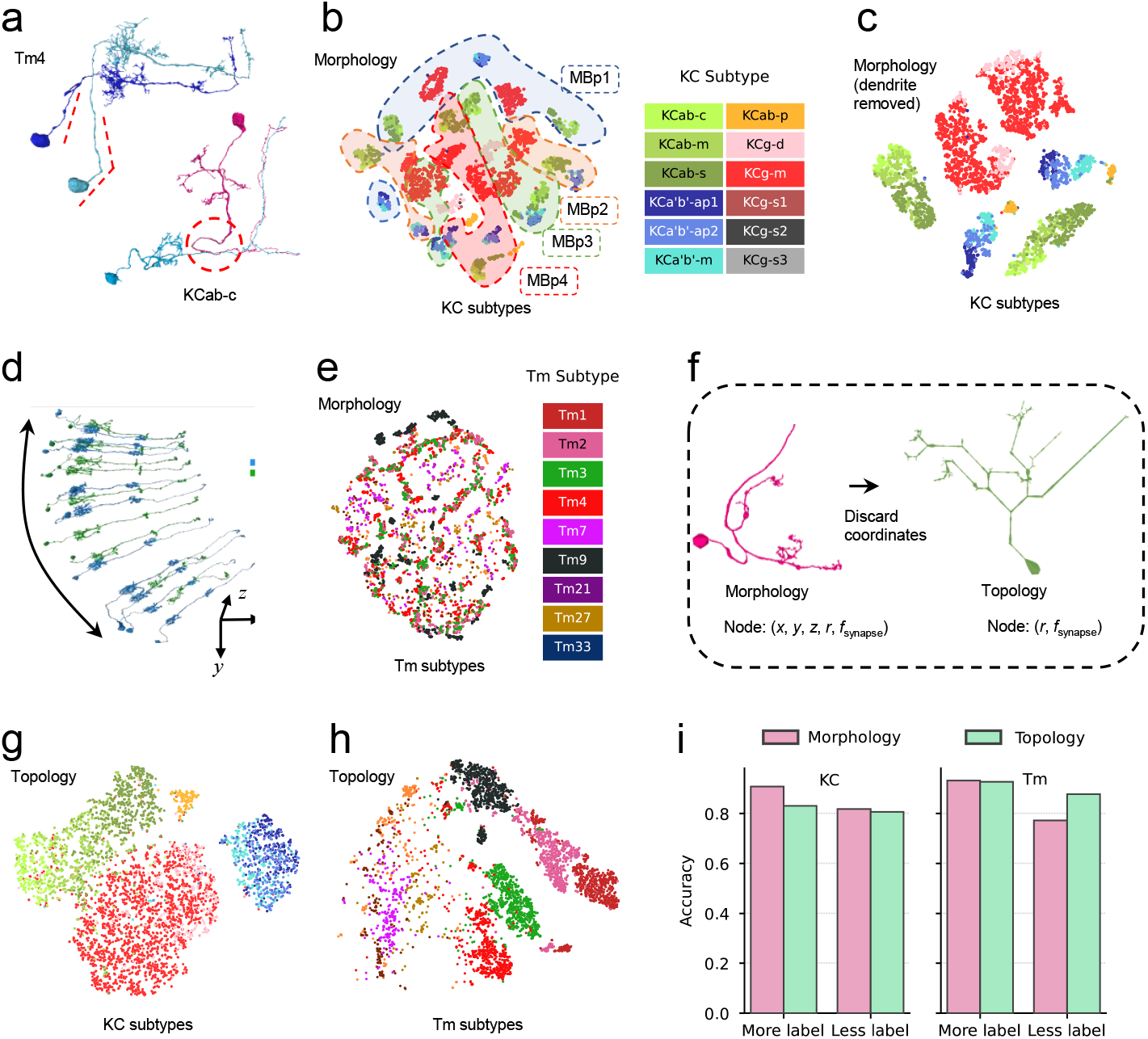
(a) Demonstration of developmental structures in Tm4 and KCab-c neurons. (b–c) t-SNE visualization of KC morphology embeddings before and after removing dendritic segments including the primary neurite, showing a substantial reorganization of the embedding space after removal of developmental structures. (d) Morphological variation of Tm neurons along the y-axis, indicating strong alignment between embedding structure and global brain geometry, where closely related types (Tm3 and Tm4) may become less separable than intra-class variability. (g-h) t-SNE visualization of topology representations of KC (g) and Tm (h) neurons. (i) Bar plots showing classification accuracy for KC and Tm tasks with topology and morphology inputs under 80% and 10% label conditions. For KC, the accuracy gap between modalities narrows under few labels; for Tm, topology outperforms morphology in the less-label setting.

Using KC and Tm neurons as examples, we first consider developmental processes that introduce non-functional morphology. In Drosophila, the proximal neurite segment between the soma and the first major branching point (often referred to as the primary neurite or pre-branch segment) largely reflects developmental elongation rather than circuit-specific computation [20, 21]. When this segment is removed, the association between embeddings and developmental lineage is substantially reduced (Fig. 2b,c, lineage-based silhouette score 0.358→ − 0.013). Tm neurons in the visual system further exhibit strong rotational symmetry as well as stereotyped developmental morphology (Fig. 2a,d), both of which are unrelated to functional identity. Consistently, their t-SNE embeddings display fragmented subtype organization (Fig. 2e, subtype silhouette score −0.073).

To disentangle structural identity from coordinate-dependent variation, we construct a topological representation that preserves node-level features while removing spatial coordinates (Fig. 2f). The resulting transformation from morphological to topological representations substantially reduces correlations between embeddings and spatial axes, indicating diminished reliance on explicit spatial encoding. Topological representations reduce confounds arising from hemispheric and developmental structure in KCs (Fig. 2g, silhouette score, hemisphere: *r* : 0.798 → −0.030; lineage silhouette: 0.358 → 0.056). For Tm, a substantial reduction in the linear correlation between the embeddings and the y-coordinate, indicating reduced reliance on explicit spatial encoding (maximum absolute Pearson’s *r*: 0.720 → 0.133; mean |*r*|: 0.274 → 0.044), while improving subtype separability in Tm neurons (Fig. 2h, silhouette score: −0.0797 → 0.020).

### Structure-driven connectivity improves subtype discrimination without partner labels

The skeleton framework also has a natural path toward connectivity analysis. Once a neuron-level embedding has been learned, that embedding can be reassigned to its associated synaptic nodes (Fig. 3a), providing a bridge between cell-level structural representation and connectivity representation. We term this paradigm structure-driven connectivity.

**Figure 3.**
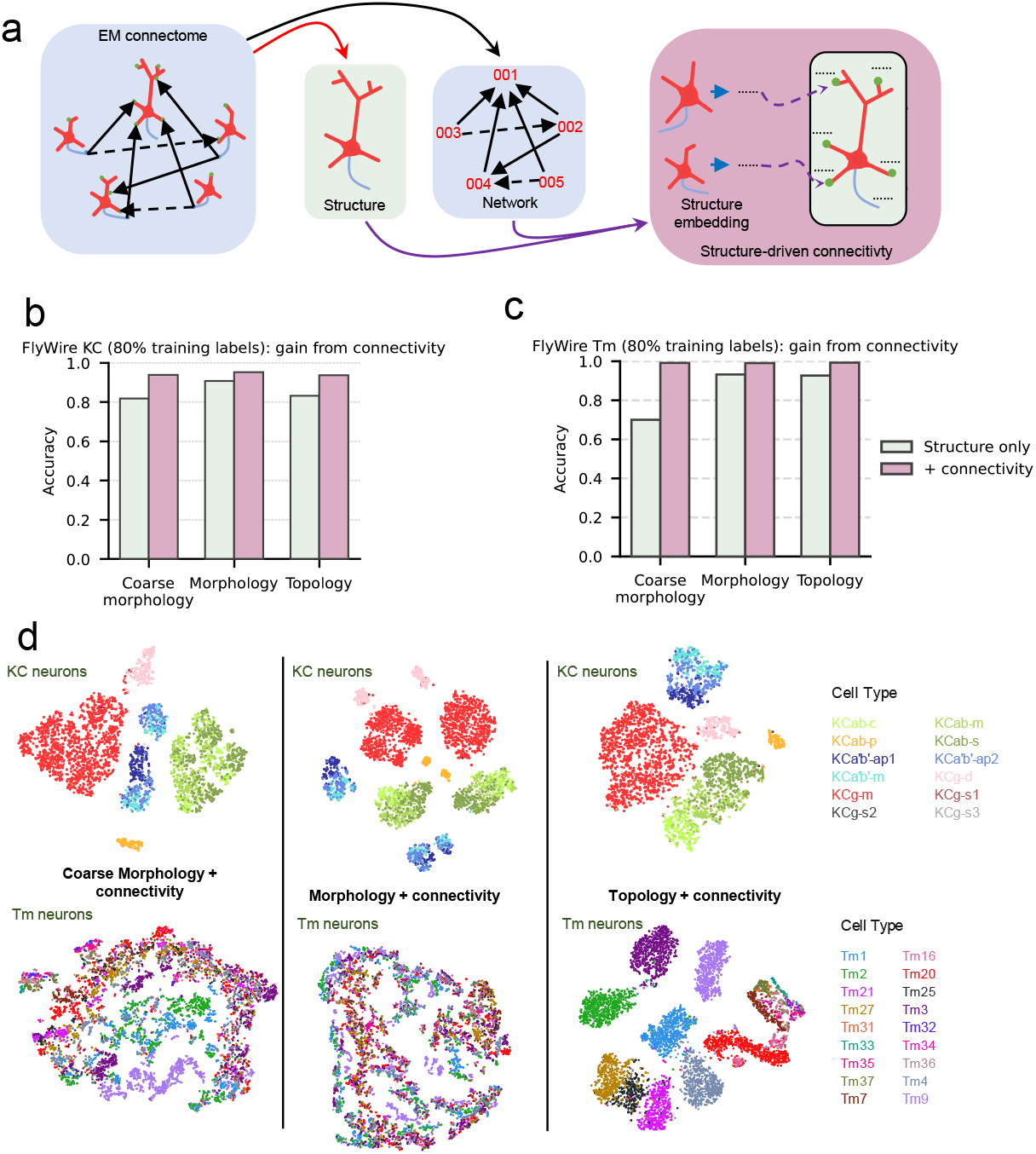
Structure-driven connectivity and classification evaluation. (a) Schematic of structure-driven connectivity, integrating fine-grained morphology and connectivity from the microscale connectome. A Transformer with graph positional encoding aggregates neighboring neurons’ embeddings. Representations are constructed from coarse morphology, fine morphology, and neuronal topology. The resulting models under coarse morphology, morphology and topology embedding are evaluated on KC (b) and Tm (c) subtype classification. Structure-driven connectivity improves KC and Tm classification accuracy across all input modalities. (d) t-SNE of topology-driven representations learned via contrastive learning from coarse morphology, morphology, and topology embeddings for KC (top) and Tm (bottom). KC subtypes cluster by topology with no hemispheric bias, while Tm neurons show clearer cluster separation with topology than with morphology-based connectivity.

We find that structure-driven connectivity consistently enhances subtype discrimination across both KC and Tm tasks (Fig. 3b,c), corroborating that synaptic context provides complementary information beyond neuronal structure alone. For both lineages, the largest gain is observed for coarse morphology (KC: +12.0 pp, 81.7%→93.8%; Tm: +29.1 pp, 70.0%→99.2%), indicating that connectivity contributes substantial subtype-discriminative information beyond intrinsic structure. Fine morphology attains the highest absolute accuracy for KC (95.2% with connectivity; 90.8% structure-only). For Tm, all three connectivity-augmented embeddings exceed 99% (topology: 92.7%→99.3%; morphology: 93.2%→99.1%; coarse: 70.0%→99.2%), showing that structure-driven connectivity is already sufficient for near-ceiling Tm subtype discrimination.

The strong classification performance suggests that structure-driven connectivity provides a sufficiently informative basis for large-scale representation learning. Without introducing manual intervention such as predefined partner-type labels into connectivity analysis, we added a projection head and a contrastive objective to the structure-driven connectivity encoder. Training on the full FlyWire dataset yielded per-neuron connectivity embeddings optimized to discriminate each cell from all others according to its local connectivity neighborhood, thereby encoding population-scale synaptic context. Among them, topology-driven connectivity embeddings show leading performance in subsequent contrastive learning and unsupervised clustering analyses ([quantitative metric TBD], Fig. 3d), suggesting strong potential for subtype discovery.

Together, these results establish the feasibility and effectiveness of structure-driven connectivity representations. Topology-driven connectivity is particularly promising for unsupervised characterization of subtype organization; in the following section we extend this framework with multi-hop connectivity to study higher-order organization in the connectome.

### Iterative multi-hop connectivity learning reveals higher-order neuronal organization

In the initial stage, the structure-driven connectivity framework only associates each neuron with the structural embeddings of its directly connected neighboring neurons, which we define as *1-hop connectivity*. After the 1-hop connectivity representation converges, the learned neuron representation can be reassigned to its associated synaptic nodes, enabling the model to capture *2-hop connectivity* (Fig. 4a). This procedure can be iteratively repeated to progressively incorporate increasingly distant connectivity information, ultimately forming a hierarchical *network representation*. Consequently, the overall framework consists of three successive phases: Phase 1, structural representation learning; Phase 2, structure-driven connectivity representation learning; and Phase 3, iterative connectivity representation learning (Fig. 4a).

**Figure 4.**
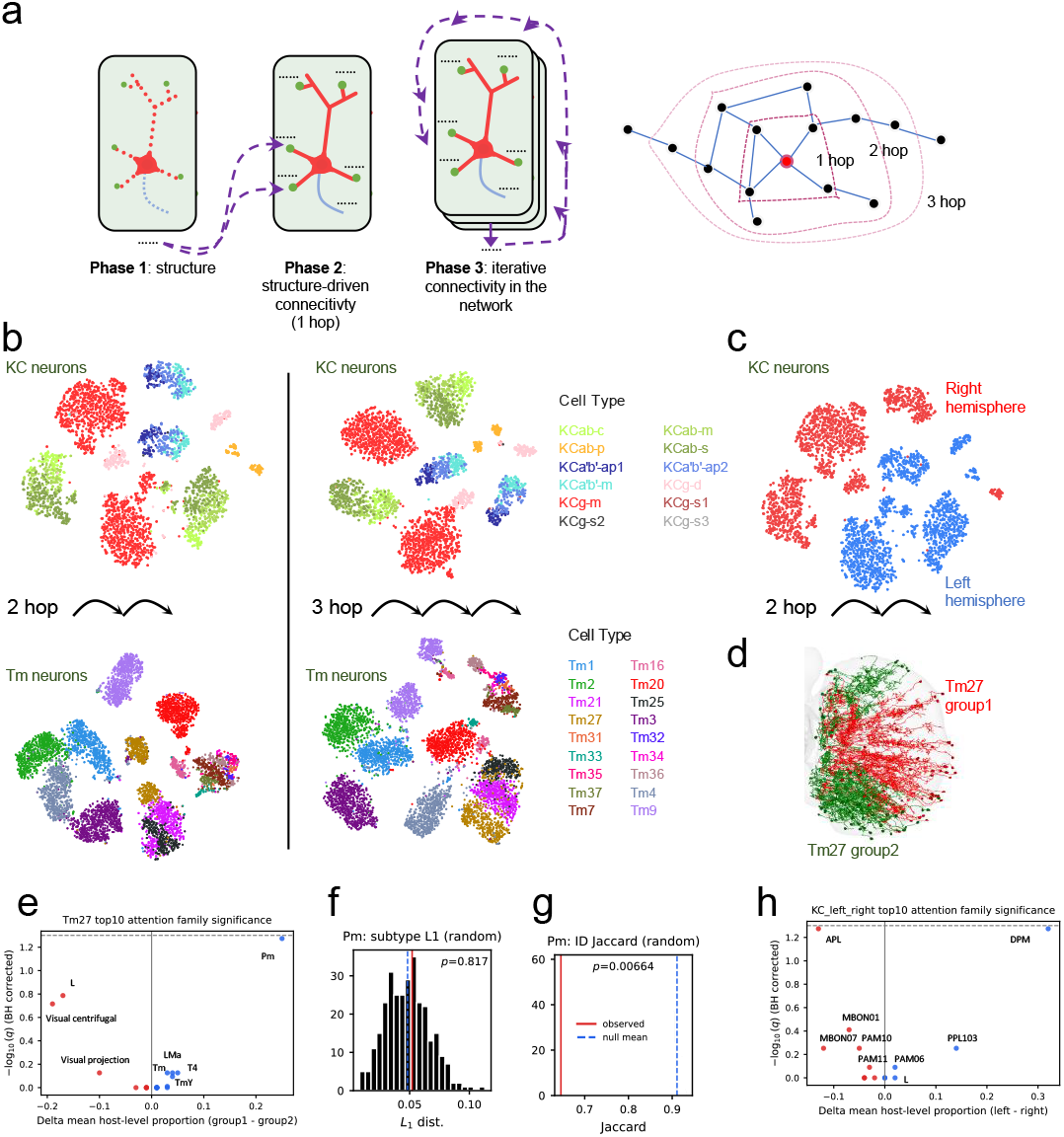
Contrastive learning of iterative connectivity representations. (a) Three phases: phase 1 – structure only; phase 2 – structure-driven connectivity; phase 3 – iterative connectivity. With more iterations, information from multi-hop neighbors is propagated to each neuron, and the iterative process converges under the deep learning and contrastive learning framework. (b) t-SNE of 2-hop and 3-hop connectivity representations, colored by KC (top) and Tm (bottom) subtypes. (c) t-SNE of 2-hop connectivity for KC colored by hemisphere. (d) Neuron renderings of two Tm27 groups. (e) High-attention connectivity partners identified by the Tm27 subgroup-classification model (finetuned on 2 hop connectivity contrastive learning model), showing that higher-order connectivity differences are primarily associated with Pm, L, and visual centrifugal neurons. (f) Quantification of the total number and proportion of Pm subtype connections in the two Tm27 groups, showing no significant difference (*p* = 0.817). (g) Comparison of connected Pm neuron populations between the two Tm27 groups, revealing significant differences in partner identity composition despite similar overall connection abundance (*p* = 0.00664). (h) High-attention connectivity partners identified by the left–right KC hemisphere classification model (fine-tuned from 2-hop connectivity contrastive learning), highlighting asymmetric connectivity patterns of APL and DPM despite their bilaterally symmetric anatomy.

Comparison between 1-hop and 2-hop connectivity representations further revealed biologically meaningful organization. Although no spatial coordinate information distinguishing the left and right hemispheres was included in the topology embeddings, KC neurons became naturally separated according to brain hemisphere in the 2-hop connectivity t-SNE visualization (Fig. 4b, c). This observation suggests that hemispheric differences in KC connectivity primarily emerge through second-hop neuronal partners rather than structural differences among direct neighbors. In contrast, the Tm27 subtype separated into two distinct clusters (DBSCAN method, silhouette score 0.42) specifically in the 2-hop connectivity representation, and this segregation was not associated with left–right hemispheric identity (Fig. 4d), indicating the presence of subtype-specific differences in higher-order connectivity organization. In the following, we further identify the specific sources underlying these connectivity differences through attention-based analysis.

To interpret the connectivity mechanisms underlying the KC and Tm27 phenomena observed in the 2-hop connectivity contrastive-learning representation, we trained binary classifiers for left–right KC classification and both Tm27 subgroup discrimination using the attention-analysis framework described in the following section, and analyzed the high-attention connectivity partners identified by the model. For Tm27, the dominant 2-hop connectivity differences were associated with connections to Pm, L, and visual centrifugal neurons (Fig. 4e). Although the total number and proportion of Pm connections did not differ significantly between the two Tm27 groups (Fig. 4f, *p* = 0.817), the specific populations of connected Pm neurons differed significantly (Fig. 4g, *p* = 0.00664), indicating that higher-order connectivity specificity rather than connection abundance distinguishes the two groups.

In the KC hemisphere-classification model, attention was primarily concentrated on connections involving APL, DPM, and a subset of MBON neurons (Fig. 4h). Among them, APL is associated with global inhibitory feedback and sparsity control in KC, whereas DPM is involved in recurrent modulatory and memory-related processes [1, 22, 23]. Despite the relatively small number of these neurons and their largely symmetric bilateral anatomical distribution, the model consistently identified asymmetric inter-hemispheric connectivity patterns, suggesting a degree of lateralized organization in mushroom body circuitry.

Leveraging iterative self-supervised contrastive learning and clustering, we further perform unsupervised connectivity-defined subgroup discovery across all known neuron types in FlyWire (Supplementary Tab. 1). This large-scale analysis identifies 84 neuron types exhibiting left–right hemispheric connectivity differences and 19 types showing higher-order connectivity organization. For both categories, only neuron types with a silhouette score > 0.5 were included. Neuron types exhibiting hemisphere-related clustering may also contain unilateral higher-order connectivity organization, which was not further disentangled here.

Importantly, across multiple hop iterations, clustering enters a convergent regime: from 3-hop to 5-hop, the number of annotated cell types supporting multi-cluster solutions changes only modestly (111→113→118; <7%, silhouette score > 0.2), whereas mean silhouette remains elevated (0.51→0.58→0.55; +42% versus structure-only topology, 0.39 at 0-hop). As each hop defines structure-driven connectivity over a distinct synaptic neighbourhood scale, identical cluster assignments across hops are neither expected nor required (Supplementary Fig.1). Results across different hop levels are not expected to be fully identical, as each hop captures connectivity organization at a distinct scale. In practical applications, fine morphology, topology, and connectivity representations across all hop levels can jointly provide complementary information for the analysis and exploration of previously uncharacterized connectomes.

Together, these results establish iterative connectome representation learning as a unified framework for integrating neuronal structure, synaptic connectivity, and higher-order circuit organization. By propagating structural representations through multi-hop connectivity in a self-supervised manner, the model captures increasingly global organizational features beyond direct synaptic neighborhoods. The emergence of consistent subtype- and hemisphere-specific patterns, together with the stability of clustering across iterations, demonstrates that connectivity-driven representations encode robust latent circuit structure. Overall, this framework provides a scalable and annotation-free approach for uncovering hierarchical organization principles in large-scale neural circuits.

### Attention localizes objective-dependent synaptic and partner features

Visualization of attention intensity on neuronal skeletons reveals distinct spatial patterns of connectivity salience under contrastive learning, KC subtype classification and KC hemisphere classification objectives (Fig. 5a–c). Which is to say, synapse-level attention is reallocated in a task-dependent manner, concentrating on connections that are most informative for the training objective. This suggests that attention can serve as an interpretable readout of connectivity differences associated with neuronal discrimination.

**Figure 5.**
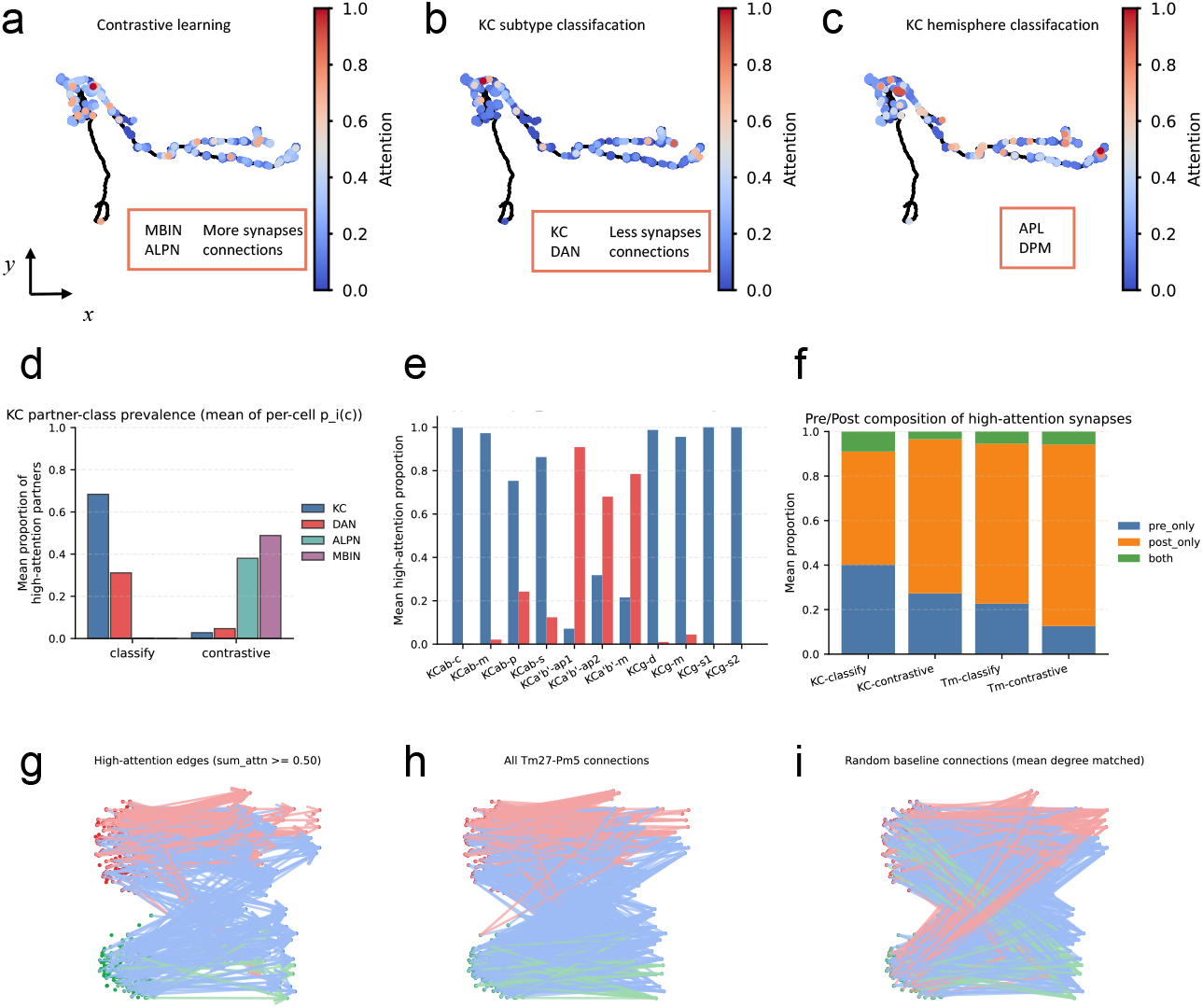
A KCg-d neuron is shown as an example, including 1-hop connectivity attention on the skeleton from a contrastive learning model (a), KC subtype classification (b), and KC hemisphere classification (c). Top attentions fall on different neighboring neurons under different models. (d) Effect of the KC subtype classification and contrastive learning models on the high attention partners. (e) Distribution of high-attention targets for KC subtypes and dopaminergic neurons (DANs), indicating subtype-specific proportions across different connectivity patterns. (f) Distribution of pre- and post-synaptic partners among high-attention synapses, showing that post-synaptic connections are more highly represented in connectivity attention. (g-i) The high-attention partners(g), real connections(h) and random connections(i) between groups(top-bottom) of Tm27(left) and Pm5(right)

By further analyzing the partner identities associated with high-attention synapses, we find that, alongside the hemispheric connectivity differences involving neighbouring neurons such as APL and DPM, attention under KC subtype classification is primarily concentrated on KC and DAN partner neurons (Fig. 5d, e). In contrast, under the contrastive learning setting, attention is predominantly assigned to ALPN and MBIN targets (Fig. 5d). These findings are broadly consistent with the known organization of the mushroom body circuit, in which KC subtype identity is closely associated with compartment-specific dopaminergic innervation[20], whereas broader KC identity is shaped by characteristic interactions with projection neurons and mushroom body input pathways[24, 25].

At the level of FlyWire partner classes among high-attention synapses, KC subtype labels (*n* = 11 subtypes represented in the exported cohort) did not yield statistically significant heterogeneity in Kenyon-cell versus dopaminergic (DAN) target proportions under the supervised classifier (Kruskal–Wallis; Kenyon-cell *P* = 0.12; DAN *P* = 0.12; Fig. 5c). Thus, within our thresholded partner-resolution assay, subtype-specific differences in DAN versus KC targeting were not resolved after classification training—consistent with subtype distinctions instead arising from *which* Kenyon or DAN identities are contacted rather than aggregate class fractions.

Across both subtype classification and contrastive learning settings, connectivity attention is consistently dominated by post-synaptic sites in both KC and Tm neurons. This suggests that upstream partner representations constitute a primary and stable substrate of structure-driven connectivity representations. (Fig. 5f).

The training attention shifts are accompanied by changes in synaptic roles. In KC, supervision increases the fraction of doubly anchored sites (0.101 vs 0.028; *P* = 0.0039), whereas in Tm it reduces post-synaptic dominance (0.754 vs 0.856; *P* = 0.023). Together, these results indicate that training objectives reorganize attention across synaptic configurations, reflecting distinct connectivity cues used for subtype discrimination.

Extended analyses that pool partner identities at root-neuron resolution (Supplementary tables; optional panels for union KC∪DAN partner overlap) can be used to test whether subtype separation operates through shared versus distinct Kenyon/DAN partners; sample sizes per subtype under contrastive training remain modest and should be interpreted cautiously.

## Discussion

We developed a scalable framework for representation learning in dense connectomes by combining hierarchical skeleton decomposition with graph contrastive learning, addressing both the structural complexity of high-density neuronal skeletons and enabling connectivity analysis driven by structural representations. The study reveals three connected principles. First, structural resolution matters: finely sampled skeletons preserve identity-relevant information that is lost under coarse skeletonization, motivating scalable graph learning directly on high-resolution neuronal trees. Second, representation geometry matters: coordinate-aware morphology is informative but can be confounded by developmental variation, spatial position and anatomical symmetry, whereas coordinate-free topology provides a more robust representation layer for clustering and label-efficient neuron typing. Third, connectivity integration matters: learned structural embeddings can serve as continuous descriptors of synaptic partners, enabling structure-driven connectivity representations without predefined partner-type labels. Together, these components form a scalable strategy for down-stream connectome interpretation with reduced dependence on curated labels and expert-defined priors.

The first principle concerns structural resolution. Across multiple FlyWire populations, finely sampled skeletons substantially improved classification compared with coarse skeletons. These gains indicate that high-resolution skeletons do not merely provide a more detailed rendering of neuronal shape; they preserve local branching patterns, synapse placement and neurite-level organization that are informative for neuronal identity. To fully exploit this information, computational strategies must also scale to the size and complexity of neuronal trees. Under current hardware constraints, HGNNA makes this practical by decomposing large neuronal trees into subgraphs, encoding local structure, and recursively integrating these embeddings into whole-neuron representations. Thus, the benefit of high-resolution skeletons depends both on the information preserved at the input stage and on the computational framework available to learn from it.

The second principle concerns representation geometry. Coordinate-aware morphology captures spatial embedding and fine anatomical detail, but it is also sensitive to developmental variation, absolute position and coordinate symmetry. These sources of variation can be biologically meaningful, as in lineage-related KC structure, but they can also dominate the latent space when the goal is to recover a coordinate-invariant notion of neuronal identity. Coordinate-free topology mitigates these effects by retaining branching organization and synapse-related node information while removing absolute coordinates, yielding a more robust representation for clustering and low-label inference. This does not imply that coordinates are unimportant; rather, morphology and topology should be treated as complementary regimes, with geometry preserved when spatial organization is the question and suppressed when it acts as nuisance variation.

The third principle concerns connectivity integration. Neuronal identity is defined not only by intrinsic structure, but also by synaptic partners. Conventional connectivity analysis often depends on predefined neuron types or expert-guided partner labels [7, 17], which limits scalability to newly reconstructed or incompletely annotated datasets. Our structure-driven connectivity representation instead uses learned structural embeddings as continuous descriptors of synaptic partners: each synaptic site on a target neuron receives the embedding of its partner neuron, and a graph neural network aggregates these partner embeddings across the target skeleton. This formulation integrates intrinsic structure with synaptic-partner context within a common representation-learning framework, improving neuronal identity discrimination beyond structural representations alone while avoiding early discretization into pseudo-types or curated partner categories. The results show that connectivity-enhanced representations improve neuronal identity discrimination beyond their corresponding structural baselines, indicating that synaptic-partner context contributes discriminative signal complementary to intrinsic morphology or topology. Contrastive connectivity learning further suggests that these representations capture a distinct layer of organization beyond structural topology alone, although effects on latent-space metrics vary across neuronal populations. Thus, connectivity does not simply sharpen all clusters uniformly, but reorganizes representation geometry in a population- and objective-dependent manner.

The attention analyses provide an initial view of what the structure-driven connectivity model learns. Supervised classification and contrastive learning reallocated synapse-level attention in different ways, suggesting that the model does not simply assign uniform salience to dense connectivity regions but learns objective-dependent patterns of synaptic importance. In KCs, supervised subtype classification concentrated attention into fewer high-salience synapses than contrastive learning, whereas Tm neurons showed the opposite tendency. Partner-class analyses further suggested that different training objectives emphasize distinct connectivity channels, with contrastive learning preserving broader ALPN/MBIN-related heterogeneity and supervised KC classification emphasizing features more directly associated with subtype discrimination. These results should be interpreted as model-based evidence of learned synaptic salience rather than direct biological mechanism. Nevertheless, they demonstrate that the framework can support interpretable downstream analyses by linking learned connectivity representations back to synaptic locations and partner classes.

Our framework supports annotation-light downstream analysis of dense connectomes by reducing reliance on curated labels and predefined partner categories. Fine skeleton learning captures high-resolution structural signal; coordinate-free topology improves robustness to geometric confounds; and structure-driven connectivity extends neuronal embeddings to synaptic-partner context. Several limitations remain. First, although topology-based representations proved robust and effective for driving connectivity learning, they remain fundamentally structural descriptions of neurons. Future work may benefit from incorporating additional modalities, such as transcriptomic, functional or developmental representations, which may provide more informative descriptors for initiating connectivity-based learning. Second, although the present framework provides separate structural and multi-hop connectivity representations, how to effectively integrate these complementary views for joint clustering and analysis remains an open challenge. Third, attention-based interpretation should also be treated cautiously and validated by perturbation, ablation or stability analyses. Finally, broader validation across additional connectomes and neuronal populations will be important for establishing generality. Despite these limitations, our results show that learned structural embeddings can serve not only as compact representations of individual neurons, but also as continuous descriptors of synaptic partners, enabling scalable analysis of neuronal structure and connectivity in dense connectomes.

**Supplementary Figure 1.**
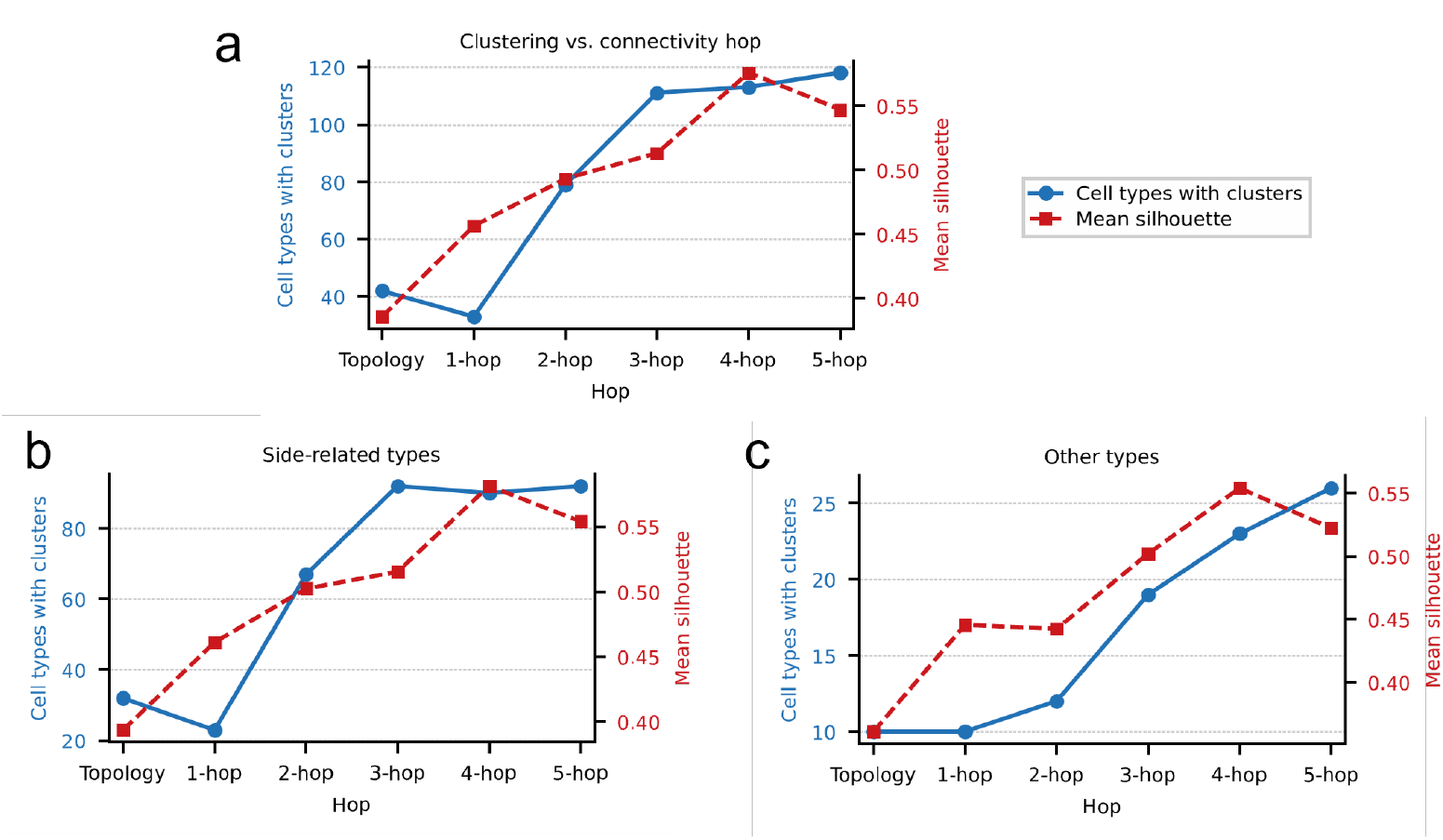
(a–c) Number of cell-type clusters discovered across multi-hop iterations. (a) Total clusters; (b) clusters associated with hemisphere; (c) clusters not associated with hemisphere.

## Method

### Datasets

We used “FlyWire Brain Dataset”, a whole-brain connectomics dataset for *Drosophila*. FlyWire artefacts processed in our pipelines include `classification.csv.gz` (cell taxonomy spreadsheet linking root identifiers to categorical annotations), `flywire_skeleton_swcs.tar.gz` (distributed morphology skeleton SWCs aligned with FlyWire v630), and `flywire_synapses_783.feather` (synapse coordinates released alongside FlyWire v783). Morphological skeleton records were obtained from the FlyWire v630 release, synaptic annotations were obtained from the corresponding FlyWire v783 synapse table, and global FlyWire-wide labels were obtained from the v630 cell-type table referenced by `classification.csv.gz`. Additional optic/visual subsets used `visual_neuron_types.csv` from the FlyWire v630 distribution.

The taxonomy spreadsheet released as `classification.csv.gz` exposes nested categorical fields (`super_class`, `class`, `sub_class`, `cell_type`, `hemibrain_type`). For Kenyon cells, rows carry family-level membership via `class`=`Kenyon_Cell`, whereas subtype-resolution identifiers such as KC subtype labels arise from finer-grained fields (`cell_type`, `hemibrain_type`). For optic transmedullary neurons (Tm), identities primarily reside under `super_class`=`optic`, with subtype distinctions indexed chiefly through `cell_type` (e.g., `Tm9`). To avoid lexical ambiguity with FlyWire’s coarse `class` column, throughout supervised KC/Tm experiments we reserve the word “class” to denote supervised subtype-resolution targets (cardinality |*C*|), independent of FlyWire’s spreadsheet column named `class`.

For Kenyon-cell (KC) subtype experiments, we used 12 classes:

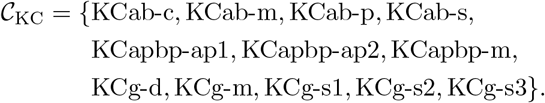

For Tm subtype experiments, we used 26 classes defined by the official FlyWire annotation:

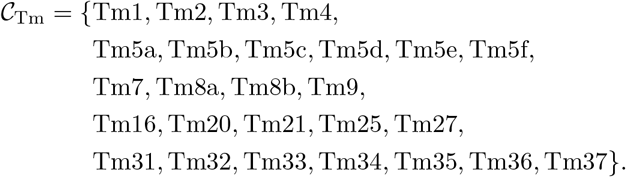

For clarity of visualization, Tm5 and Tm8 were omitted from the t-SNE plot in Fig.2 to Fig.5 and are presented separately in the Supplementary Figure.

### Fine skeleton construction

#### Nodes

Each neuron skeleton was represented as a graph *G* = (*V, E*) loaded from per-neuron skeleton records. Each node initially stored geometric and semantic attributes derived from SWC-like morphology and synaptic annotations, including coordinates (*x, y, z*), radius *r*, soma flag, and synaptic role features.

#### Skeleton

Node indices were remapped to contiguous integers for stable batching. For morphology and topology processing, skeletons were represented in a soma-centered coordinate system so that the soma sits at the origin. Skeletons exceeding the predefined limit (3500 nodes) were subsampled while protecting key nodes (e.g., soma and semantically protected nodes).

#### Morphology, topology, and connectivity features

##### Morphology/topology branch

In morphology-oriented settings, geometric channels (e.g., coordinates and derived edge descriptors) were retained. In topology-only settings, raw coordinates were removed and models used compact topological/synaptic descriptors. Depending on the experiment, Fourier positional coding and geometric augmentations were enabled.

##### Connectivity branch

For neuron-connectivity models, only synapse-related nodes were retained:

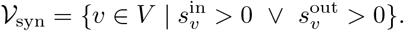

Node features were replaced by precomputed topology embeddings:

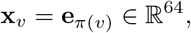

where *π*(*v*) is the embedding index stored in the node feature table (last column), and **e** is a pretrained topology embedding. For training-time regularization, synaptic-node dropout—random subsampling of synaptic nodes with a prescribed retention probability—and Gaussian embedding noise were applied:

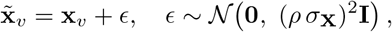

with *ρ* the embedding-noise rate and *σ*_**X**_ the sample standard deviation of node embeddings in the current graph.

### Hierarchical skeleton decomposition

For hierarchical GraphDINO training, each neuron graph was decomposed into two levels:

1. First-level subgraphs of fixed size *s*_1_ (typically *s*_1_ = 8) by BFS-style partitioning.
2. Second-level groups of first-level subgraphs with size *s*_2_ (typically *s*_2_ = 8), yielding a compressed whole-graph representation.

At each level, adjacency matrices were constructed and Laplacian positional encodings were computed from the corresponding adjacency.

### Hierarchical graph neural network architecture

Our core encoder is a graph Transformer with attention logits formed from query-key similarity and graph adjacency priors:

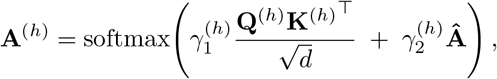

where **Â** is the adjacency matrix (with task-specific padding mask), and *γ*_1_, *γ*_2_ are learned node-wise trade-off factors. Given values **V**, attention output is

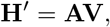

followed by residual MLP blocks.

For hierarchical GraphDINO, we used three encoders:

- node-to-subgraph encoder (*d*_sub1_ = 10, depth 5),
- subgraph-to-supergraph encoder (*d*_sub2_ = 24, depth 6),
- whole-graph encoder (*d* = 64, depth 7, 4 heads).

For connectivity-only models, we used a single graph Transformer encoder (dim 64, depth 6, 4 heads, max pooling) followed by either a projection head (contrastive) or a classification head.

### Memory-cost analysis

For a one-level decomposition of an *N* -node graph into subgraphs of size *n*, the total adjacency memory scales as

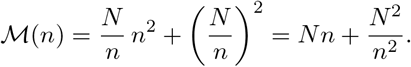

Differentiating gives

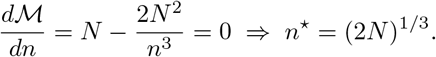

Thus decomposition converts one *N* ^2^ adjacency into a linear-plus-compressed term and substantially reduces peak memory in practice, especially when combined with synaptic-node dropout.

### Contrastive learning

For contrastive pretraining, all neurons in the global training-index set were treated as training samples. For each neuron, two stochastic graph views (*G*_1_,G_2_) were drawn independently via synaptic-node dropout and embedding-noise perturbation (detailed in the following subsection). Student and teacher encoders produced logits **z**^(*s*)^ and **z**^(*t*)^, with teacher updated by EMA:

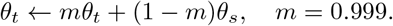

Teacher centering used EMA with decay 0.9. The cross-view DINO objective was

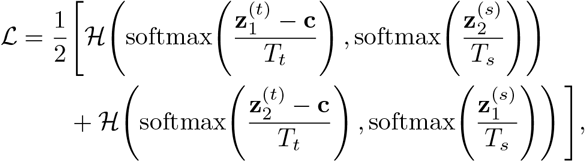

where *T*_*t*_ = 0.06, *T*_*s*_ = 0.9, and **c** is the teacher center.

### Data augmentation

Data augmentation was applied depending on the branch (morphology, topology, or connectivity) and training objective; the schemes below are not tied exclusively to supervised classification—connectivity-oriented perturbations also underpin contrastive pretraining.

#### Morphology

Morphology pipelines employed geometric perturbations—coordinate jitter, global translation, and mirroring—in addition to the topology-oriented perturbations described below. The mirror flip exploits approximate bilateral symmetry between left and right hemispheres.

#### Topology

Topology pipelines applied random node deletion, random branch pruning, and a hierarchical random graph partitioning scheme, where each neuronal skeleton was first decomposed into fixed size blocks typically via breadth first expansion before performing localized perturbations, inspired by DINOv2 in the field of computer vision [26].

#### Connectivity

Connectivity pipelines disabled geometric channels; stochastic regularization operated in embedding space via synaptic-node dropout and Gaussian embedding noise, reducing overfitting to fixed pretrained embeddings.

### Classification test and latent-space analysis

For supervised neuron subtype classification, the final classification head was trained using the cross-entropy loss:

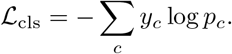

We evaluated topology-based and morphology-based classifiers using a consistent data split protocol across all neuron types. Reported classification accuracy corresponds to the best validation accuracy achieved within the first 300 training epochs. For latent-space analysis, neuron embeddings from trained models were exported and visualized in 2D using t-SNE to assess cluster separation and subtype consistency.

#### Classification accuracy evaluation metrics

Let *V* denote the validation set of size *N* = |*V*|. For each sample *i* ∈ *V*, let *y*_*i*_ ∈ {1, …, *K*} be the ground-truth class and *ŷ*_*i*_ the predicted class. The *micro-averaged* accuracy (overall accuracy) on the full validation set is

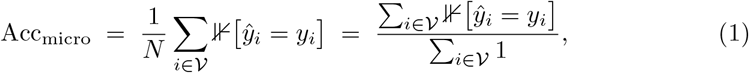

where ⊮[·] is the indicator function.

To reduce instability from classes with extremely few validation examples, we additionally report accuracy computed after *excluding all validation samples whose true class is rare*. For each class *k*, let

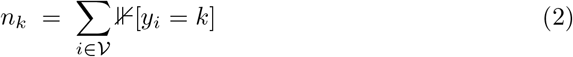

be the number of validation samples with label *k*. Given a minimum support threshold *m* ∈ ℕ (we use *m* = 2 unless stated otherwise), define the set of *eligible* (non-rare) classes

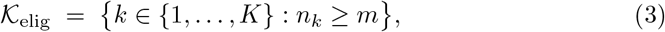

and the corresponding subset of validation indices

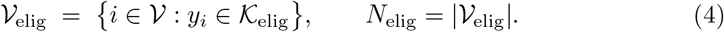

The *rare-class-excluded micro accuracy* is then the standard accuracy restricted to *V*_elig_, i.e. the same numerator–denominator rule as Acc_micro_ but with both restricted to eligible samples:

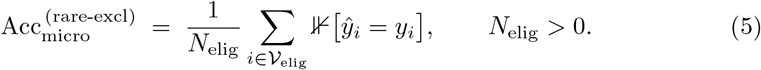

Equivalently, samples with *y*_*i*_ ∉ *K*_elig_ contribute neither to the numerator nor to the denominator. If *N*_elig_ = 0, the metric is undefined and can be reported as not applicable.

### Embedding and connectivity analysis

Structure-driven connectivity model used synapse-only induced graphs with topology-embedding node features.

### Tm27 group comparison

We fine-tuned a binary Tm27 classifier (`Tm27_group1` vs. `Tm27_group2`) from the contrastive-learning checkpoint of the 2-hop connectivity model (`ckpt_91.pt`, using an 80% training split, and inferred synapse-level attention from the epoch-13 model weights (`ckpt_13.pt`). For each group, 10 representative neurons were selected for inference. We computed synapse-level attention scores within each neuron and retained the top 10 attention-weighted partner neurons per sample. Each selected partner was further mapped to its corresponding neuron identity and annotation labels, including `visual_type`, `family_type`, and `super_class`. To account for originally unassigned family labels (`none`), we additionally evaluated two relabeling strategies: `none`→`visual_type` and `none`→`super class`.

To test whether a partner category is significantly enriched in one Tm27 group over the other, we performed host-level permutation tests. For each host neuron *i*, we computed the within-host proportion of one-hop partners belonging to category *c*:

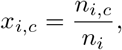

where *n*_*i,c*_ is the number of one-hop partners in category *c*, and *n*_*i*_ is the total number of one-hop partners for host *i*. Group-level means were then

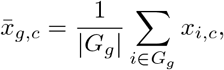

and the observed effect size was

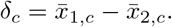

For each category *c*, we pooled host-level values {*x*_*i,c*_} from both groups and generated a null distribution of *δ*_*c*_ by random label permutation (5000 permutations, preserving group sizes). One-sided and two-sided *p*-values were computed from this null distribution:

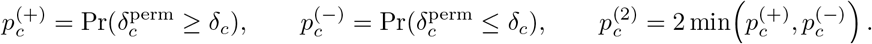

Multiple testing across categories was controlled using the Benjamini–Hochberg procedure, yielding *q*-values 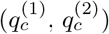. Results were visualized in an effect-size significance plot with x-axis *δ*_*c*_ and y-axis − log_10_ 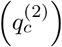; categories above the *q* = 0.05 threshold were considered statistically significant.

### Statistics of Attention

For each setting (KC/Tm *×* classify/contrastive), we used the exported table `attention_top_partners.tsv` (thresholded by attention ≥ 0.7) as synapse-level observations. Cell-level summaries were computed by grouping rows by (host_cell_type, host_neuron_id):

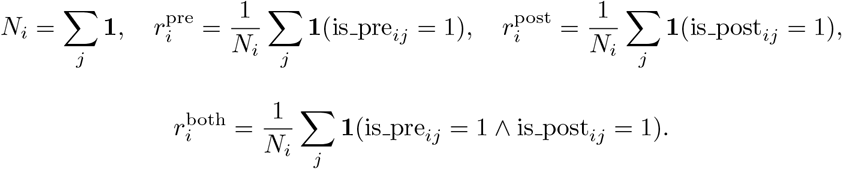

For partner class *c* ∈ {Kenyon_Cell, DAN, ALPN, MBIN}, the per-cell target proportion is

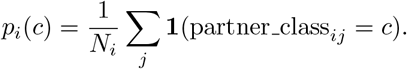

Model comparison (classify vs contrastive) within KC or Tm was performed on matched neurons (same host_neuron_id) using two-sided Wilcoxon signed-rank tests for 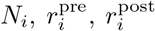, and 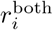. For KC subtype analysis, subtype-level differences in *p*_*i*_(*c*) were tested using Kruskal–Wallis tests across KC subtypes. All plots report distribution summaries or subtype means derived from the same cell-level tables.

### All type Cluster-number estimation across representation spaces

To quantify whether neuron subtypes exhibit distinct sub-structure in different representation spaces, we estimated cluster numbers in a unified, reproducible pipeline and compared results across *Topology, Topology-driven Connectivity*, and *2-hop Connectivity* embeddings.

#### Step 1: Cell-type-specific embedding extraction and normalization

For each target cell type, neuron IDs were collected from curated annotation tables (Kenyon-specific table when available, otherwise FlyWire classification table), excluding types with neuron counts below a predefined threshold. Matched embedding vectors were extracted and z-score normalized per feature dimension.

#### Step 2: Graph construction for spectral clustering

A *k*-nearest-neighbor graph was constructed in the standardized embedding space (nearest-neighbor affinity). This graph provides a geometry-aware substrate for non-convex cluster structure that is not well captured by centroid-only partitioning.

#### Step 3: Automatic cluster-number selection (eigengap)

Instead of scanning many fixed cluster counts, we used the eigengap criterion on the normalized graph Laplacian:

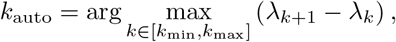

where *λ*_*k*_ are ordered Laplacian eigenvalues. This yields a data-driven cluster count for each cell type and representation.

#### Step 4: Spectral partitioning and quality readout

Given *k*_auto_, spectral clustering was run once to obtain final labels. We then computed the silhouette coefficient in Euclidean space as a compact separation/compactness summary:

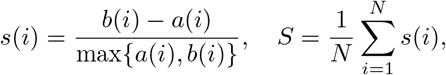

where *a*(*i*) is the mean intra-cluster distance for sample *i*, and *b*(*i*) is the minimum mean distance to other clusters.

#### Step 5: Conservative interpretive threshold (optional reporting layer)

For conservative interpretation in downstream summaries, we optionally applied a silhouette floor (*S >* 0.2) to distinguish practically meaningful partitioning from weak fragmentation. This threshold was used as a reporting criterion, not as the clustering objective itself.

**Table 1:**
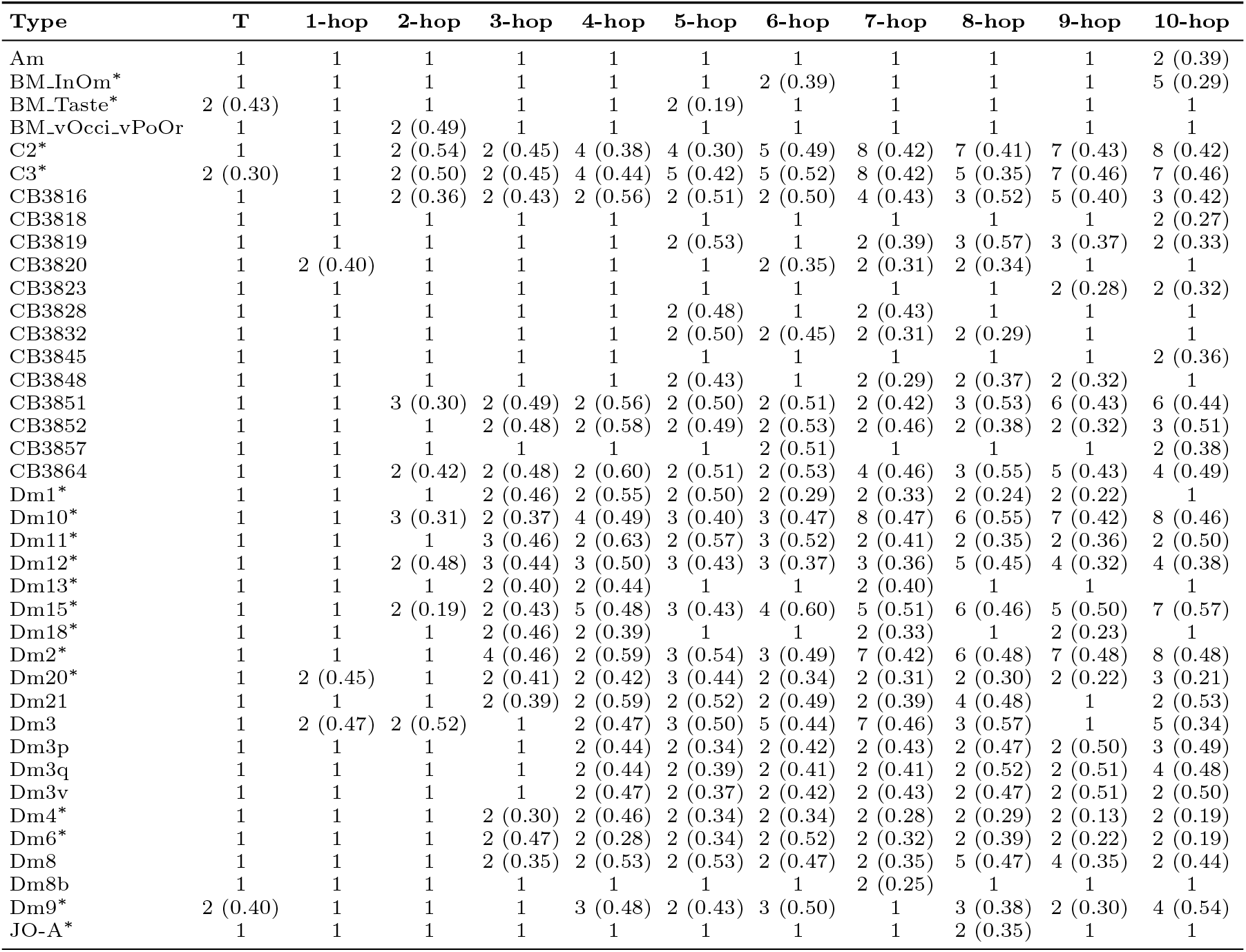
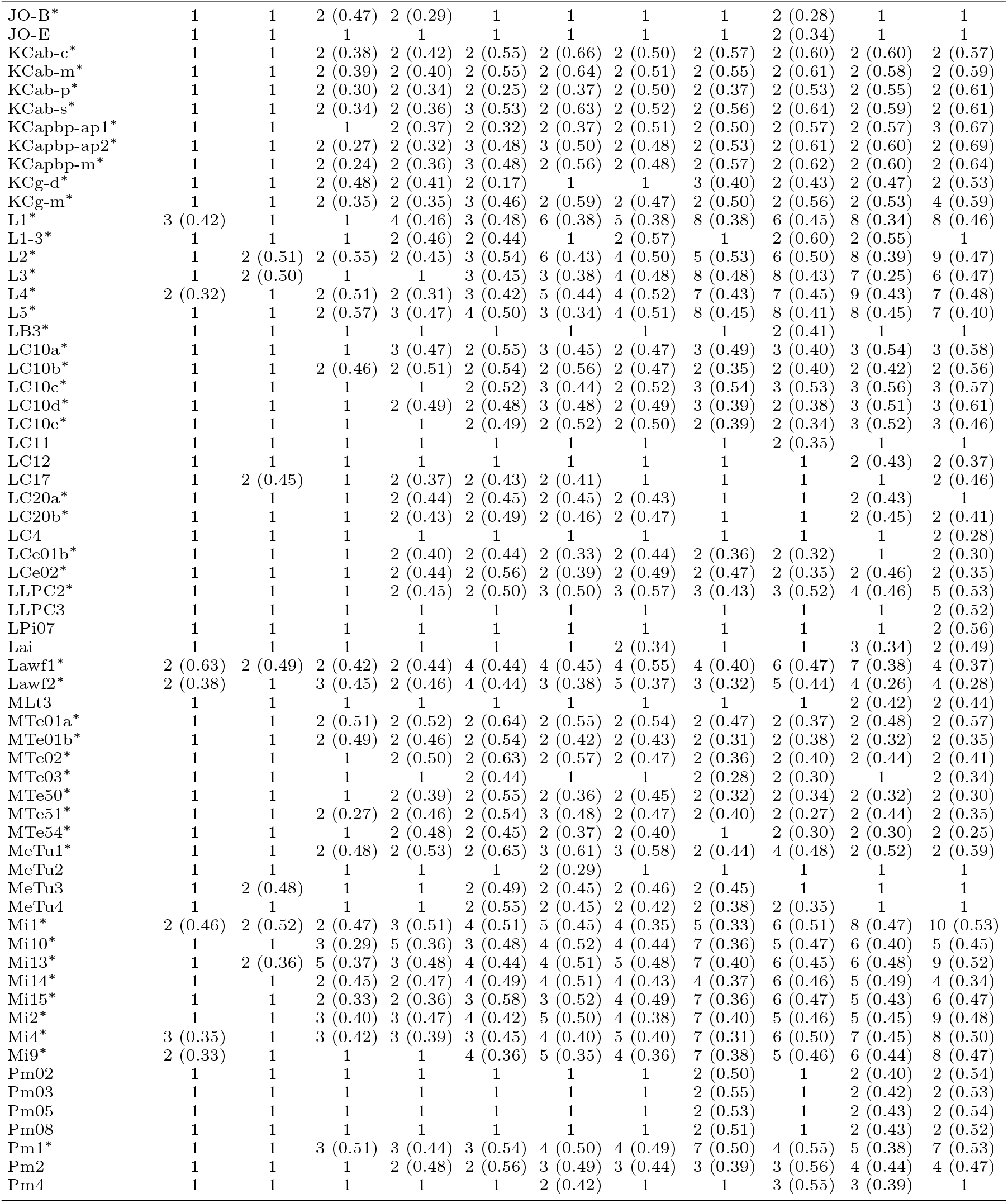
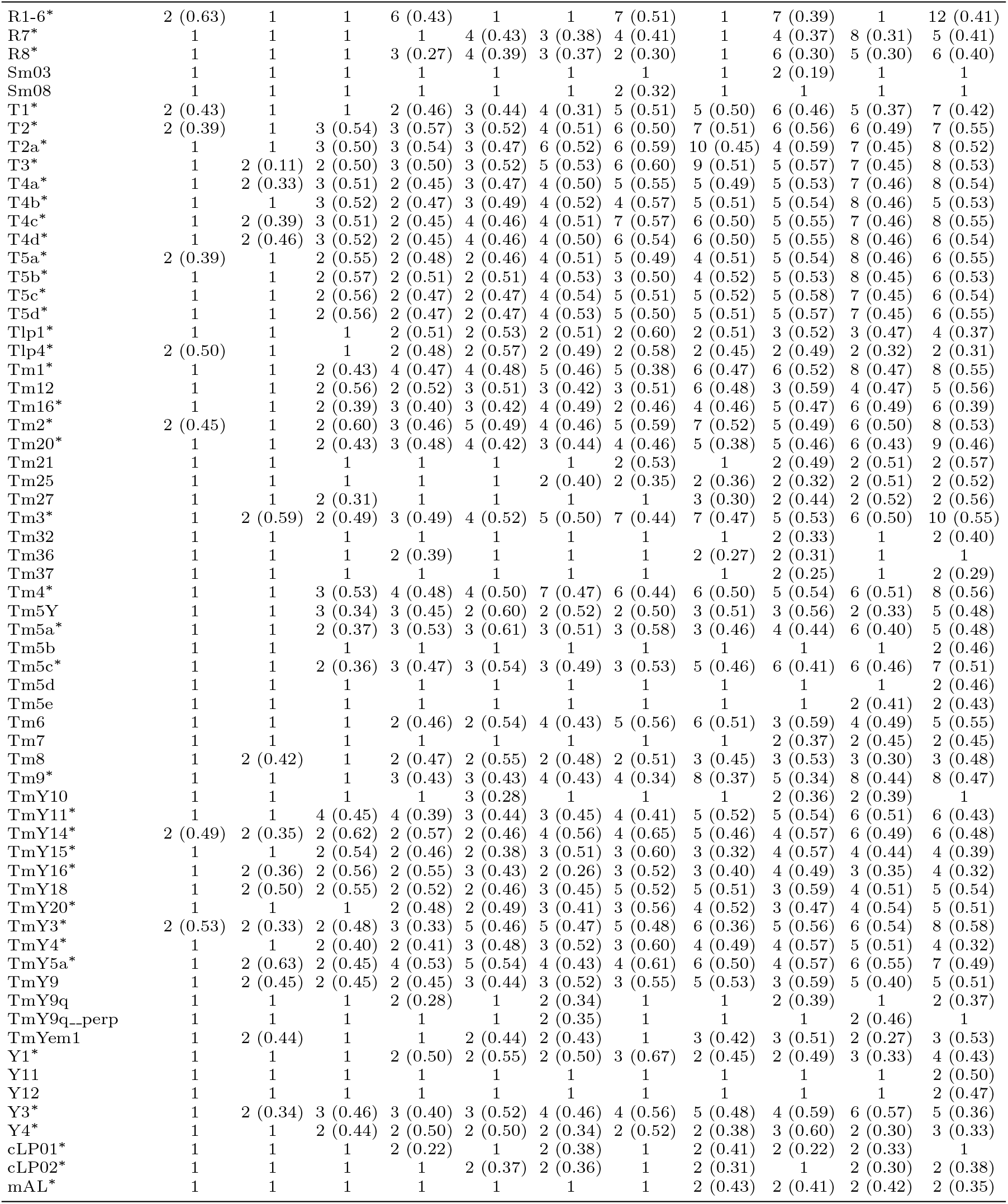
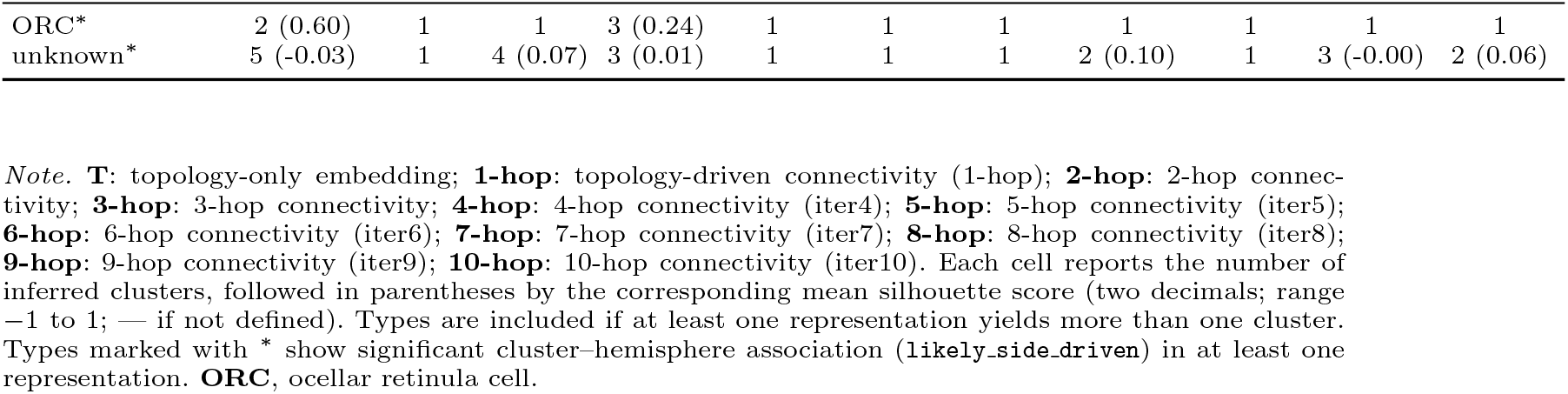
Inferred cluster counts and silhouette by representation.

## References

[1] Scheffer, L.K., et al.: A connectome and analysis of the adult Drosophila central brain. eLife 9, 57443 (2020)

[2] Dorkenwald, S., et al.: Neuronal wiring diagram of an adult brain. Nature 634, 124–138 (2024)

[3] Ding, Z., et al.: Functional connectomics reveals general wiring rule in mouse visual cortex. Nature 640, 459–469 (2025)

[4] Motta, A., et al.: Dense connectomic reconstruction in layer 4 of the somatosensory cortex. Science 366, 3134 (2019)

[5] Schneider-Mizell, C.M., et al.: Inhibitory specificity from a connectomic census of mouse visual cortex. Nature 640, 448–458 (2025)

[6] Elabbady, L., et al.: Perisomatic ultrastructure efficiently classifies cells in mouse cortex. Nature 640, 478–486 (2025)

[7] Schlegel, P., et al.: Whole-brain annotation and multi-connectome cell typing of Drosophila. Nature 634, 139–152 (2024)

[8] Ganguly, I., Heckman, E.L., Litwin-Kumar, A., Clowney, E.J., Behnia, R.: Diversity of visual inputs to Kenyon cells of the Drosophila mushroom body. Nature Communications 15, 5698 (2024)

[9] Lin, A., et al.: Network statistics of the whole-brain connectome of Drosophila. Nature 634, 153–165 (2024)

[10] Celii, B., et al.: NEURD offers automated proofreading and feature extraction for connectomics. Nature 640, 487–496 (2025)

[11] Weis, M.A., et al.: An unsupervised map of excitatory neuron dendritic morphology in the mouse visual cortex. Nature Communications 16, 3361 (2025)

[12] Schmidt, H., Gour, A., Straehle, J., Boergens, K.M., Brecht, M., Helmstaedter, M.: Axonal synapse sorting in medial entorhinal cortex. Nature 549, 469–475 (2017)

[13] Ren, H., Dai, H., Dai, Z., Yang, M., Leskovec, J., Schuurmans, D., Dai, B.: Combiner: Full attention Transformer with sparse computation cost. In: Advances in Neural Information Processing Systems (NeurIPS), vol. 34, pp. 22429–22442 (2021)

[14] Dorkenwald, S., Li, P.H., Januszewski, M., Berger, D.R., Maitin-Shepard, J., Bodor, A.L., Collman, F., Schneider-Mizell, C.M., da Costa, N.M., Lichtman, J.W., Jain, V.: Multi-layered maps of neuropil with segmentation-guided contrastive learning. Nature Methods 20, 2011–2020 (2023)

[15] Chiang, W.-L., Liu, X., Si, S., Li, Y., Bengio, S., Hsieh, C.-J.: Cluster-GCN: An efficient algorithm for training deep and large graph convolutional networks. In: Proceedings of the 25th ACM SIGKDD International Conference on Knowledge Discovery & Data Mining, pp. 257–266 (2019)

[16] Panicker, C.A., Geetha, M.: Exploring graph partitioning techniques for gnn processing on large graphs: A survey. In: 2023 4th International Conference on Communication, Computing and Industry 6.0 (C216), pp. 01–07 (2023). IEEE

[17] Matsliah, A., et al.: Neuronal parts list and wiring diagram for a visual system. Nature 634, 166–182 (2024)

[18] Schwartzman, G., Jourdan, B., García-Soriano, D., Matsliah, A.: NTAC: Neuronal type assignment from connectivity. Nature Communications 17, 1284 (2026)

[19] Bates, A.S., Manton, J.D., Jagannathan, S.R., Costa, M., Schlegel, P., Rohlfing, T., Jefferis, G.S.X.E.: The natverse, a versatile toolbox for combining and analysing neuroanatomical data. eLife 9, 53350 (2020)

[20] Li, F., et al.: The connectome of the adult Drosophila mushroom body provides insights into function. eLife 9, 62576 (2020)

[21] Costa, M., Manton, J.D., Ostrovsky, A.D., Prohaska, S., Jefferis, G.S.: Nblast: rapid, sensitive comparison of neuronal structure and construction of neuron family databases. Neuron 91(2), 293–311 (2016)

[22] Pitman, J.L., Huetteroth, W., Burke, C.J., Krashes, M.J., Lai, S.-L., Lee, T., Waddell, S.: A pair of inhibitory neurons are required to sustain labile memory in the drosophila mushroom body. Current Biology 21(10), 855–861 (2011)

[23] Liu, X., Davis, R.L.: The gabaergic anterior paired lateral neuron suppresses and is suppressed by olfactory learning. Nature neuroscience 12(1), 53–59 (2009)

[24] Zheng, Z., Lauritzen, J.S., Perlman, E., Robinson, C.G., Nichols, M., Milkie, D., Torrens, O., Price, J., Fisher, C.B., Sharifi, N., et al.: A complete electron microscopy volume of the brain of adult drosophila melanogaster. Cell 174(3), 730–743 (2018)

[25] Caron, S.J., Ruta, V., Abbott, L.F., Axel, R.: Random convergence of olfactory inputs in the drosophila mushroom body. Nature 497(7447), 113–117 (2013)

[26] Oquab, M., et al.: DINOv2: Learning robust visual features without supervision. arXiv preprint arXiv:2304.07193 (2023)

